# Tumoral Hypoxic Extracellular Vesicles Foster a Protective Microenvironment in Triple-Negative Breast Cancer

**DOI:** 10.1101/2024.11.01.621519

**Authors:** Bianca C. Pachane, Pedro H. T. Bottaro, Aline M. Machado, Cynthia A. de Castro, Gabriela Guerra, Larissa T. Gozzer, Marina M. Grigoli, Artur D. Zutião, Angelina M. Fuzer, Marcia R. Cominetti, Wanessa F. Altei, Heloisa S. Selistre-de-Araujo

## Abstract

The highly metastatic triple-negative breast cancer (TNBC) relies on the tumor microenvironment (TME) to maintain phenotypic heterogeneity and progression. Extracellular vesicles from hypoxic TNBC (EVh) have been previously shown to facilitate tumoral invasion; however, their function in the tumor microenvironment remains unclear. We used a novel method to investigate the TME *in vitro* called multicellular circulating co-culture, to characterize how EVh interferes with tumoral and endothelial cells, fibroblasts, monocytes and macrophages. EVh promoted monocyte differentiation to M2-like macrophages and inhibited macrophage-derived phagocytosis in endothelial and tumoral cells. The protection of endothelial, tumoral and stromal cellular integrity by EVh increased pro-tumoral and pro-angiogenic signaling, collagen matrix synthesis and showed a potential differentiation to cancer-associated fibroblasts. Our findings highlight the critical role of EVh in protecting tumor cells, indicating its cooperation towards a protective TME, which was demonstrated by the multicellular circulating co-culture and conventional co-culture protocols. These findings lead to an adequate system with potential for investigating other tumor-related processes, including circulating tumor cells and metastasis.

**Graphical Abstract:** 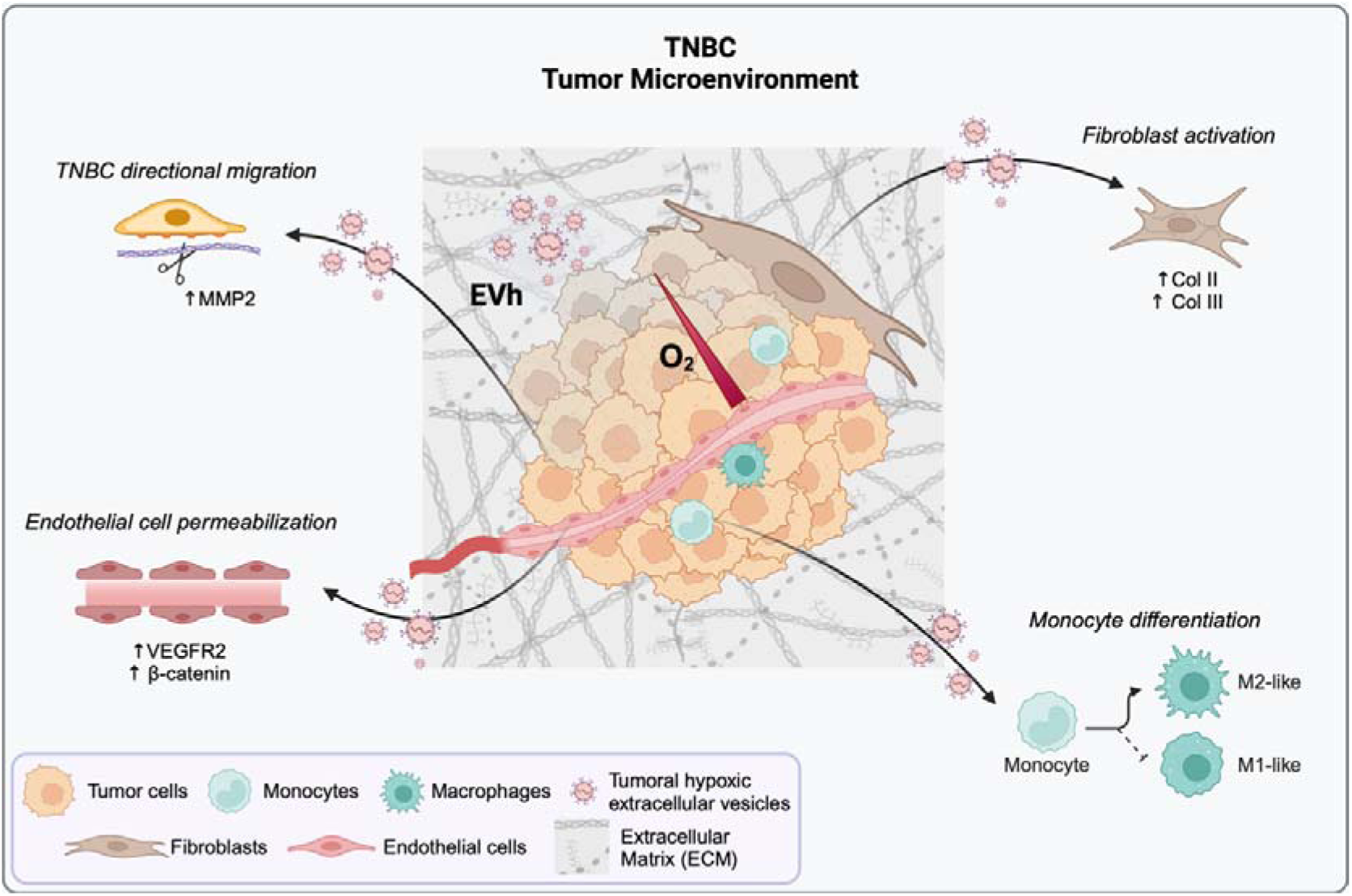

## Background

Triple-negative breast cancer (TNBC) is mainly characterized by the lack of expression of receptors for estrogen, progesterone, and the human epidermal growth factor (HER2) (1). Comprised of a heterogeneous mass with a high risk of metastasis, TNBC affects over 15% of all breast tumors and presents poor overall survival (2). Its development is plagued with hypoxia, a condition in which oxygen distribution is reduced due to the increased cellular activity within the tumoral tissue, the inefficient tumoral vasculature, and the constant competition for nutrients within the mass (3,4). Hypoxia and tumorigenesis favor the secretion of extracellular vesicles (EV), nanoparticles (30-300 nm) released by all known cells, with a major function in intercellular communication (5,6). We previously showed that tumor-derived hypoxic EVs (EVh) favor pro-invasive pathways when compared to normoxic tumoral EVs (7), but how they affect the TNBC tumor microenvironment (TME) remains unknown.

The TME is a myriad of cellular and acellular components that evolve as the tumor phenotypic heterogeneity grows according to the stages of cancer progression (8–10). In the pre-invasive TME, autocrine signaling by epithelial (EGF), hepatocyte (HGF), and transforming growth factors (TGF-β) favors cell proliferation, migration, and plasticity (11). The extracellular matrix (ECM) is primarily modified by fibroblasts, specifically cancer-associated fibroblasts (CAF), through protein synthesis and the expression of remodeling enzymes, which alters the protein content, activity, and crosslinking to favor collagen synthesis and deposition (12,13). Pro-inflammatory mediators induce monocyte differentiation into macrophages and their subsequent polarization into tumor-associated macrophages (TAM), whose phenotype may vary from the classically polarized, pro- inflammatory M1 subtype to the alternatively polarized, anti-inflammatory M2 phenotype (13). The increased cell activity of the tumor stimulates the release of reactive oxygen species, which favors DNA damage and the secretion of pro-angiogenic stimuli, including VEGF and fibroblast growth factor (FGF-2) (11). Once the invasive profile is achieved, cell motility is favored by the polarization of the cytoskeleton, the interplay between adhesion molecules, and ECM remodeling by matrix metalloproteases (MMPs) (14). These changes in cell behavior result from the secretion of TGF-β, interleukins (IL-6 and IL-8), and VEGF by CAFs, which favors the epithelial-mesenchymal transition, survival, and maintenance of immune control (11).

The collective complexity of the TME renders its study challenging, particularly when using *in vitro* techniques such as 2D cell culture. Due to the intrinsic method limitations, including the restriction in the number of simultaneous cell cultures and the low correspondence with the physiology of clinical TME, there has been a shift towards 3D cell culture models, which add a novel layer of complexity to *in vitro* studies. However, one pitfall of these models is the difficulty in observing individual cellular responses from the mass (15). In this work, we describe a novel method for studying the TME *in vitro* called the multicellular circulating co-culture (MC-CC), to determine how TNBC-derived hypoxic EVs influence individual immortalized cellular models in a TME setting. Using this approach, we demonstrated for the first time that EVh protects cells from macrophage-derived phagocytosis. This pro-tumorigenic function of EVh confirms previous suppositions on the tumoral function of EVs and supports the development of novel EV-based strategies to overcome resistance to anticancer therapies.

## Materials and Methods

### Cell culture

Human cells from triple-negative mammary adenocarcinoma MDA-MB-231 (ATCC^®^ CRM-HTB-26^™^) were grown between passages 36 and 45 in Leibovitz L-15 media supplemented with 10% FBS (pH 7.0, Vitrocell) at 37 °C. Dermal fibroblasts HDFa (ATCC^®^ PCS-201-012^™^) and umbilical cord endothelial cells HUVEC (ATCC^®^ CRL-1730^™^) were maintained between passages 6 and 15 in DMEM media supplemented with 10% FBS and 1% pen/strep (pH 7.4, Vitrocell) at 37 °C, 5% CO_2_. Monocytes THP-1 (ATCC^®^ TIB-202^™^) were kept under suspension in RPMI 1640 media with added 10% FBS and 1% pen/strep (pH 7.0, Vitrocell) at 37 °C, 5% CO_2_. Cells were grown under sterile conditions for no longer than 4 weeks after thawing and were frequently tested for *Mycoplasma* using the MycoAlert^®^ Assay kit (Lonza). For co-culture, cells were maintained in OptiMEM I Reduced Serum Media supplemented with 1% pen/strep (Gibco).

### EVh separation and characterization

MDA-MB-231 cells (3000 cells/cm^2^) were seeded in EV-depleted media (antibiotic-free DMEM supplemented with 10% UC-FBS, 4.6 g/L glucose and 1 mM sodium pyruvate) and maintained for 24 h at a hypoxic atmosphere (1% O_2_, 5% CO_2_, 37 °C; H35 Hypoxystation, Don Whitley Scientific). The media was replaced by antibiotic-free OptiMEM for 48 hours, and cells were further maintained for 48 hours in hypoxia. Hypoxic EVs (EVh) were separated by differential ultracentrifugation and filtration, as described previously (7). Following the MISEV2023 guidelines (16), EV preparations were characterized by nanoparticle tracking analysis, transmission electron microscopy, protein quantification and Western blotting.

### Nanoparticle Tracking Analysis (NTA)

EVh samples were diluted in PBS (1:1000, v/v) and injected into the module of a Nanosight NS300 (Malvern). Capture settings were set at camera level 16, slide shutter 1300, slider gain 512, FPS 25.0, and 749 frames, acquired in three-60s events. Analysis was performed on NTA software (v. 3.2 Dev Build 3.2.16) using the detection threshold of 4.

### Protein quantification

EVh samples were lysed with 2% SDS, diluted in deionized water (4:150) and added to a 96-well plate with an albumin standard curve with known values (0-16 µg.ml). Using the microBCA kit (Thermo-Fisher Scientific), the working solution was prepared (25A:24B:1C) and added to samples, which were incubated for 2 hours at 37 °C. Plates were evaluated using a SpectraMax i3 plate reader (Molecular Devices) at O.D._562nm_ and quantification was considered successful if R^2^ < 0.99.

### Western blotting

MDA-MB-231 EVh (10 µl) and cell (10 µg) lysates were mixed with Laemmli buffer under reducing conditions (1:4, v/v) and boiled at 100 °C for 5 minutes. Precision Plus Protein Dual-Color (Bio-Rad) was used as the loading control. Samples were added to 10% SDS-PAGE gels for electrophoresis at 100 V and proteins were transferred to nitrocellulose membranes (0.45 µm, Bio-Rad) at 100 V, 4 °C, for 2.5 hours in buffer (2.5 mM Tris, 2 M glycine, 20% methanol; pH 7.6). Transfer efficiency was confirmed by Ponceau S membrane staining. Membranes were blocked with 5% milk-TBST for 1 hour under agitation and probed overnight for ALIX (1:1000, 186429, Abcam), Calnexin (1:1000, mAb 2679, Cell Signaling), CD63 (1:1000, 59479, Abcam) or Flotillin-1 (1:1000, 61020, BD Biosciences).

After three 5-minute washes with TBST, membranes were incubated with respective secondary antibodies (Goat anti-Rb, 1:15.000, #205718, Abcam; or Goat anti-Ms, 1:10.000, #97040, Abcam) for 1 hour at RT under agitation. Membranes were washed four times with PBST for 5 minutes and exposed to ECL substrates (Clarity^™^ Western ECL Substrate, Bio-Rad and SuperSignal^™^ West Femto, Thermo-Fisher Scientific) for 2 minutes. The chemiluminescence reaction was documented using ChemiDoc^™^ XRS^+^ (Bio-Rad).

### Transmission Electron Microscopy (TEM)

An EVh dilution (1:2, v/v in PBS) was deposited in copper grids covered with formvar and carbon (Lot 051115, 01800 F/C, 200 mesh Cu, Ted Pella Inc.) for 20 min, RT. Samples were fixed in 2% paraformaldehyde (PFA-PBS) for 20 min at RT and washed extensively in deionized water. Grids were exposed to 4% Uranyl-acetate (pH 4) and 2% methylcellulose solution for 10 min on ice and in the dark. Samples were air-dried and visualized in an FEI TECNAI G2 F20 HRTEM microscope at 40,000x magnification.

### Monocyte differentiation in macrophages

THP-1 cells were seeded in 6-well plates (6x10^5^ cells/ml) in OptiMEM medium supplemented with 1% pen/strep and treated with the vehicle control (i.e., PBS), EVh (10^9^ particles/ml) or phorbol-12-myristate-13-acetate (PMA, 100 nM), following a previously established protocol (17). Cells were incubated for 72 h at 37 °C, 5% CO_2_, and differentiation to macrophages was determined by the adhesion of cells to the well, followed by phenotype modification, assessed by light microscopy at 20x (Lux 2, CytoSMART).

### Cell viability assay

At the end of cellular assays, a 10% resazurin solution (0.1 ml/ml, #199303, Sigma-Aldrich) was prepared in OptiMEM and replaced the conditioned media of each well for 4 h at 37 °C, 5% CO_2_. A blank no-cell control and a 100%-reduced control, obtained from the reduction of resazurin by heat, were maintained in parallel. The supernatant was transferred to a new plate and analyzed by fluorescence (λ_Exc_ 555 nm, λ_Em_ 585 nm) using a plate reader (SpectraMax i3, Molecular Devices). Cell viability was determined after normalization by the blank no-cell control and displayed as the % of reduced resazurin.

### Time-lapse cell motility assay

In a 24-well dish, MDA-MB-231 (5x10^4^ cells/ml), HUVEC (5x10^4^ cells/ml) or HDFa (10^4^ cells/ml) were seeded in complete growth media (Leibovitz L-15 or DMEM supplemented with 10% FBS and 1% pen/strep) for adhesion over 24 h at 37 °C, 5% CO_2_, followed by media replacement with OptiMEM 1% pen/strep. THP-1 cells were diluted in OptiMEM 1% pen/strep to achieve a 10^5^ cells/ml concentration. Cells were treated with EVh (10^9^ particles/ml) for 24h at 37 °C, 5% CO_2_ and kept in parallel with an untreated control (i.e., PBS). Motility was evaluated under time-lapse light microscopy (Lux2, CytoSMART), with snaps taken every 30 minutes, and single-cell tracking was processed using the *TrackMate* plugin in FIJI (18,19).

### Enzyme-Linked Immunosorbent Assay - ELISA

To evaluate the levels of cytokines IL-6, IL-10, TNF-α and IL-1β from the monocyte differentiation assay and the multicellular co-culture assay, we used high-affinity microplates sensitized with monoclonal anti-cytokine antibodies (IL-6: 555220; IL-10: 555157; TNF-α: 555212; IL-1β: 557953, OptEIA, BD Biosciences) for 16-18 h. Plates were washed thrice (0.05% Tween 20-PBS), incubated in 4% FBS-PBS for 1 hour at RT, and washed again. The conditioned media and a standard curve of recombinant cytokines were added to the wells for 2 h, RT, and the excess was washed off. Next, the anti-cytokine antibodies conjugated with peroxidase were incubated for 1.5 h (RT) followed by five washes with 0.05% Tween 20. A developing solution (3,3’,5,5’-tetramethylbenzidine, TMB) was added to wells, and the reaction was stopped with sulfuric acid. Plates were screened at O.D._450nm_ (SpectraMax i3, Molecular Devices) and sample concentrations were calculated from the titration curve of the cytokine standards, expressed in pg/ml.

### Fluorescent gelatin invasion assay

Black 96-well plates were coated with fluorescein-conjugated gelatin from pig skin (0.2 mg/ml, G13187, Molecular Probes) before cell seeding. MDA-MB-231, HUVEC, HDFa, or THP-1 cells in their respective culture media were added to the gelatin-coated wells and incubated overnight at 37 °C, 5% CO_2_. The media was replaced with OptiMEM 1% pen/strep containing EVh (10^9^ particles/ml) or PBS for 24 h at 37 °C, 5% CO_2_. After incubation, cell viability was assessed using a resazurin-based fluorimetric assay, while the area of gelatin degradation and cell morphology metrics were evaluated on FIJI (ImageJ) using epifluorescence microscopy images, as previously described (7). The full protocol is available on the protocols.io platform (20).

### Cell invasion in transwell co-culture

To evaluate cell invasion in co-culture without contact between cell lines, fluorescein-labelled gelatin-coated coverslips were prepared as described previously (7) and pre- conditioned in 24-well plates with Leibovitz L-15 medium for 30 min at 37 °C before the seeding of MDA-MB-231 cells (5 x 10^4^ cells/ml). In inserts with 0.3 µm membranes, HUVEC (5x10^4^ cells/ml) or HDFa (10^4^ cells/ml) cells were seeded on the upper chamber in DMEM 10% FBS. Inserts and coverslips were incubated for 24 hours at 37 °C, 5% CO_2_ for cell adhesion. Once the monolayers were stable, a Boyden’s chamber was assembled to contain the gelatin-seeded MDA-MB-231 in the bottom chamber and an insert in the upper chamber. The system was maintained in OptiMEM for 24h at 37 °C, 5% CO_2_. Cells and conditioned media were processed following the full protocol described elsewhere (23). Downstream analysis included the evaluation of cell morphology metrics, tumoral invasion into fluorescent gelatin and matrix metalloprotease secretion by zymography.

### Gelatin zymography

To assess MMP-2 and MMP-9 activity, conditioned media samples (10 µg) were mixed with non-reducing Laemmli buffer (2:1, v/v) and loaded in technical duplicates into 10% gelatin (100 µg/ml) SDS-PAGE gels with the Precision Plus™ Dual-Color Protein Ladder (Bio-Rad) as loading control. A normalizing FBS sample (10 µg) was added to all gels for band quantification. Electrophoresis was run at 4 °C under 80 V. Gels were washed with 2.5% Triton X-100 for 40 min, RT, and submersed in refolding buffer (20 mM Tris, 5 mM CaCl_2_, 1 μM ZnCl_2_, pH 8.0) for 20 h at 37 °C. Gels were stained with Coomassie Brilliant Blue overnight, de-stained for band revelation, and photographed in ChemiDoc^™^ XRS^+^ (Bio-Rad) (22). Band densitometry was performed on FIJI (ImageJ).

### Gelatin invasion in direct co-culture

MDA-MB-231 (5x10^4^ cells/ml), HUVEC (5x10^4^ cells/ml), HDFa (10^4^ cells/ml), or THP-1 (10^5^ cells/ml) cells were stained either with CellTracker™ Red CMTPX (5 µM, C34552, Invitrogen) or CellTrace™ CSFE (5 µM, C34554, Invitrogen) in OptiMEM 1% pen/strep following the product datasheets. A choice of two cell lineages was simultaneously seeded into wells of a fluorescein-conjugated gelatin-coated 96-well black plate. A control group (PBS) was maintained alongside an EVh-treated group (10^9^ particles/ml) at 37 °C, 5% CO_2_ for 24 h. After incubation, cell viability was assessed using a resazurin-based fluorometric assay and the area of gelatin degradation and cell morphology metrics were evaluated on FIJI (ImageJ) using epifluorescence microscopy images, as previously described (7). The full protocol is available on the protocols.io platform (21).

### Multicellular circulating co-culture (MC-CC)

MDA-MB-231 (5x10^4^ cells/ml), HUVEC (5x10^4^ cells/ml) and HDFa (10^4^ cells/ml) cells were seeded on gelatin, Matrigel, and FN-coated coverslips, respectively, overnight at 37 °C, 5% CO_2._ After adhesion, they were individually assembled in sterile QV500 (*Kirkstall*) culture chambers with 1 ml of OptiMEM 1% pen/strep. THP-1 cells (10^5^ cells/ml) were pre- stained with CellTracker™ Red CMTPX (5 µM, C34552, Invitrogen) and transferred to the reservoir containing EVh (10^9^ particles/ml) or vehicle (PBS). The tubing system was closed and connected to the MDA-MB-231, HUVEC, and HDFa chambers, respectively, and propelled by a peristaltic pump set to a 50 µl/s flow rate. The circuit was incubated at 37 °C, 5% CO_2_ for 24h. Coverslips were individually processed for cell staining and immunofluorescence, following the full protocol published elsewhere (24). The conditioned media was collected, spun a 300 *x g* for 10 min, and the supernatant was probed for cytokines via ELISA and MMPs by gelatin zymography.

### Individual static and circulating culture controls

Cells seeded in ECM-coated coverslips (24) were assembled individually in a sterile QV500 (*Kirkstall*) culture chamber with 1 ml of OptiMEM 1% pen/strep and connected with the tubing system to a reservoir containing 13 ml of media. Following the previous method, the system was maintained with a peristaltic pump, parallel to a static individual control. Coverslips were removed from the compartments, counterstained, and imaged at 40x magnification (ImageXpress Micro XRS, Molecular Devices). Cell circularity index and gelatin degradation were quantified using FIJI (7,18).

### Statistical analysis

Cellular assays were repeated on three distinct occasions in technical triplicates. MC-CC was repeated four times with both groups running simultaneously. Each sample was applied twice on zymograms, and average band densitometry was used for statistics. Datasets were checked for outliers using ROUT’s test and distribution by Shapiro-Wilk (n < 9) or D’Agostino-Pearson omnibus K2 (n ≥ 9). Parametric data was evaluated using unpaired t-tests (2 groups) or ANOVA one-way with Tukey’s multiple comparison test (3+ groups). Non-parametric data was analyzed using the Mann-Whitney test (2 groups) or Kruskal-Wallis analysis of variance with Dunn’s multiple comparison test (3+ groups). Values of p < 0.05 were considered statistically relevant (GraphPad Prism (v. 9.3)).

## Results

### EVh characterization

In this follow-up study, we maintained the isolation of EVh samples protocol published before, where samples were thoroughly characterized by label-free proteomics in comparison with their normoxic counterparts (7). Incorporating the recommendations outlined in the MISEV2023 guidelines (16), EVh samples were separated from 2.2 × 10^7^ MDA-MB-231 cells grown in hypoxia in serum-free media, leading to a population with modal size of 130.2 nm (± 78.7) and concentration of 3.69 × 10^11^ particles/ml (± 8.74 × 10^10^) (Fig. 1A). Samples contained 236.6 µg/ml of protein and yielded 167,346.94 particles per cell or 1.07 × 10^-5^ µg protein per cell. Known EV biomarkers CD63, ALIX and flotillin-1 were abundant in EV samples, whereas calnexin was absent (Fig. 1B). Visualization by TEM confirmed EVh abundance and typical morphology (Fig. 1C).

**Figure 1.**
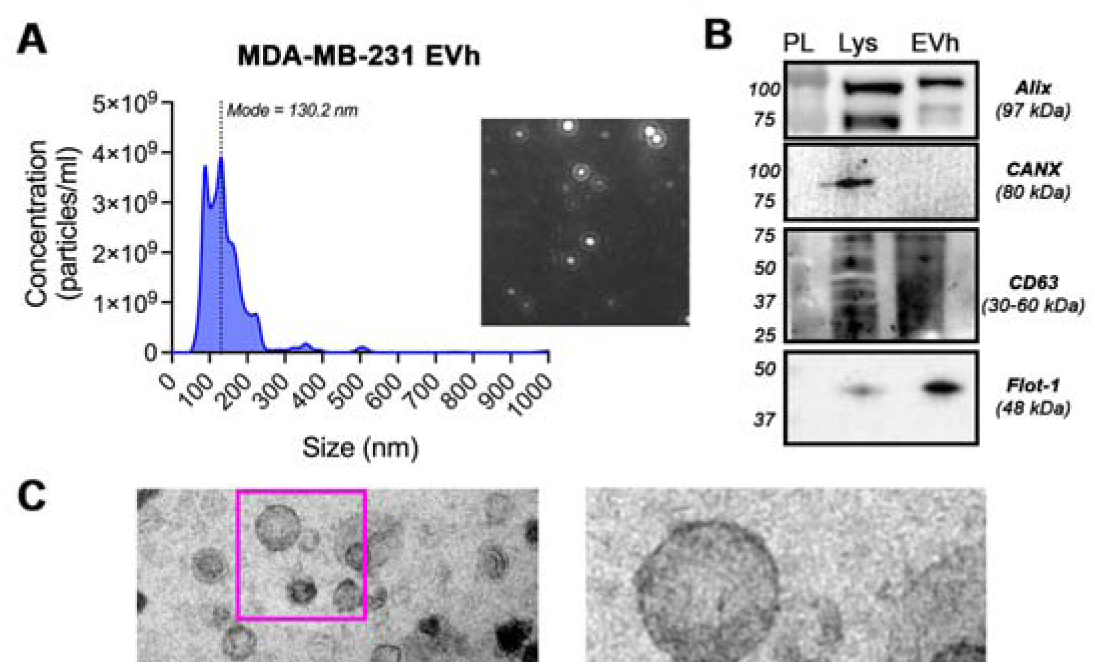
EVh characterization. (A) Size dispersion and EV abundance by nanoparticle tracking analysis of MDA-MB-231-derived EVh. (B) Detection of ALIX, calnexin (CANX), CD63 and flotillin-1. (C) Wide-field and close-up TEM images of EVh. Scale bars: 200 nm and 100 nm.

### EVh-mediated THP-1 differentiation into M2-like macrophages

To evaluate whether EVh affected the homeostatic conditions of cells found in the TME, we designed individual experiments to assess behaviors like macrophage differentiation, cell viability, invasion and motility. Using the THP-1 monocytic cell line, which grows in suspension and acquires an adherent and broader phenotype upon differentiation into macrophages, we observed that both EVh- and PMA-treated groups became adherent after 16-24 hours of treatment, while untreated cells remained in suspension for up to 72 h (Fig. 2A and Suppl. 1A,B). In contrast, while the full well of PMA-treated cells differentiated into unpolarized macrophages, a reduced population of monocytes exposed to EVh fulfilled the differentiation, which indicates a distinct differentiation mechanism between groups (Fig. 2B).

**Figure 2.**
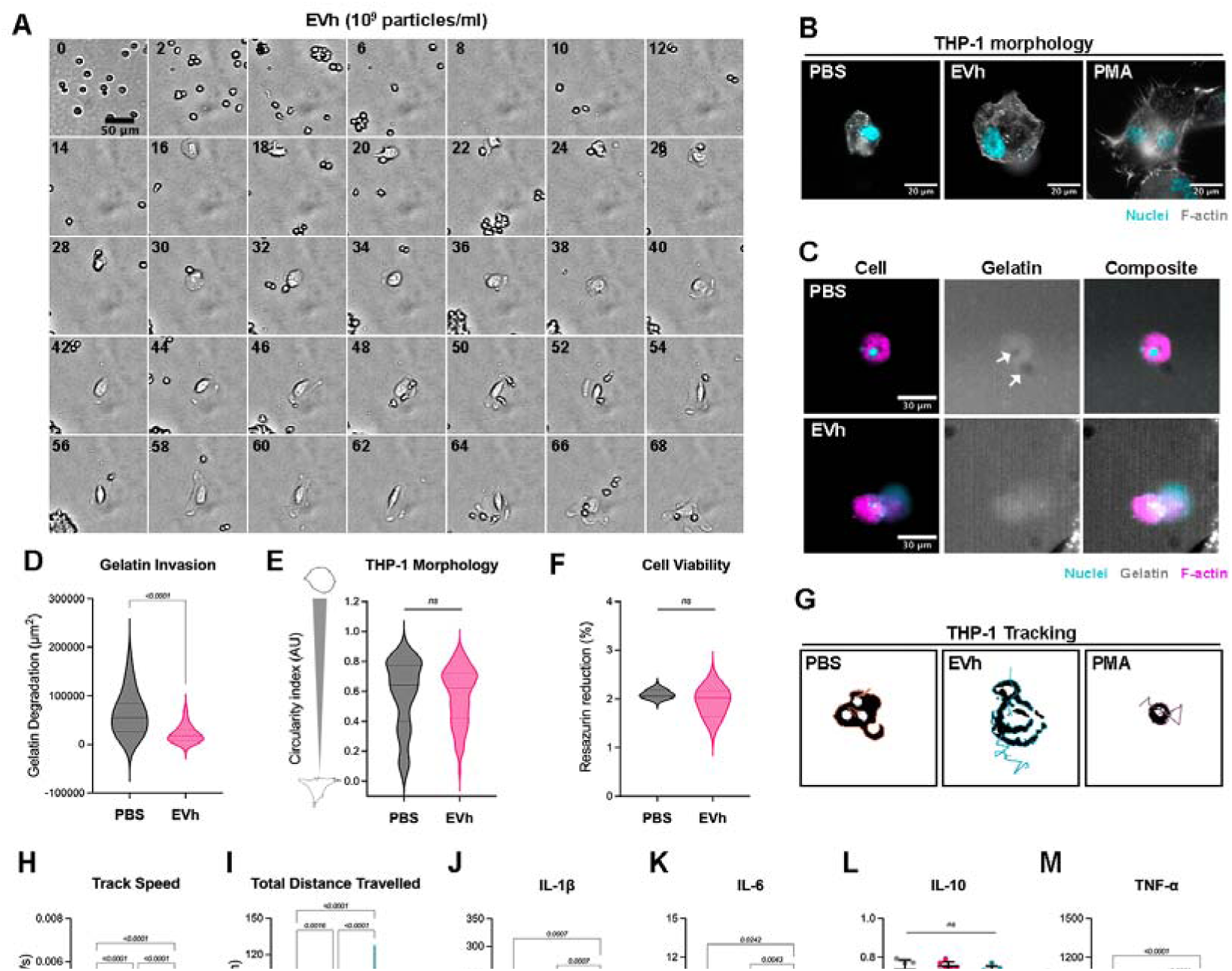
EVh-mediated macrophage differentiation. **A -** Timelapse tracking of THP-1 cells treated with EVh. Snaps from the same site were compiled in 2-hour increments over 3 days (scale bar: 50 µm). **B –** Representative images of THP-1 treated with vehicle (PBS), EVh or PMA after 72h differentiation (nuclei in cyan and F-actin in gray; scale bar: 20 µm). **C –** Representative images of THP-1 cells (nuclei in cyan, F-actin in magenta) treated with PBS or EVh on a gelatin coating (gray), with degradation spots identified by white arrows (scale bar: 30 µm). **D –** Quantification of degraded gelatin area (µm^2^). **E -** Cell circularity index for quantification of THP-1 morphology. **F -** Cell viability of cells under invasion assay. **G –** Tracking of THP-1 trajectory over 24h. **H –** Average track speed of cells in 24 h (µm/s). **I -** Total distance traveled by cells in 24 h (µm). **J-M -** Quantification of cytokines TNF-α, IL-1β, IL-6, and IL-10, respectively. Assays were repeated over three different occasions with technical duplicates or triplicates. *p* values indicated above comparative bars with statistical significance.

The EVh-induced group showed a reduction in gelatin invasion compared to the undifferentiated control (Fig. 2C,D), but no changes were observed regarding cell morphology metrics and viability (Fig. 2E,F). Cell tracking revealed undirected movement in both untreated and PMA-differentiated cells, while EVh-differentiated cells exhibited directional migration with higher mean speed and greater average distance traveled (Fig. 2G-I). Inflammatory cytokine profiling of THP-1 conditioned media detected TNF-α and IL-1β only in the PMA group (Fig. 2J,K), a non-significant trend of IL-6 reduction in the EVh group (Fig. 2L), and continuity of IL-10 expression in all three groups (Fig. 2M). Overall, these results suggest the macrophage-like cells derived from EVh signaling display a pro-tumoral behavior, which is indicative of an M2-like phenotype.

### Tumoral reprogramming of endothelial cells in vitro to promote vascular permeability

Regarding the endothelial cell model HUVEC, EVh promoted gelatin invasion and elongated cell morphology metric, which is consistent with a pro-invasive phenotype (Fig. 3A-C). Despite the clear indication of cell polarization, cell viability remained unaffected (Fig. Suppl. 2A). Unlike THP-1, there was non-directional motility in EVh-exposed HUVEC (Fig. 3D). Although the average speed of movement was similar between the groups, a higher distance traveled by cells was seen in EVh groups (Fig. 3E,F).

**Figure 3.**
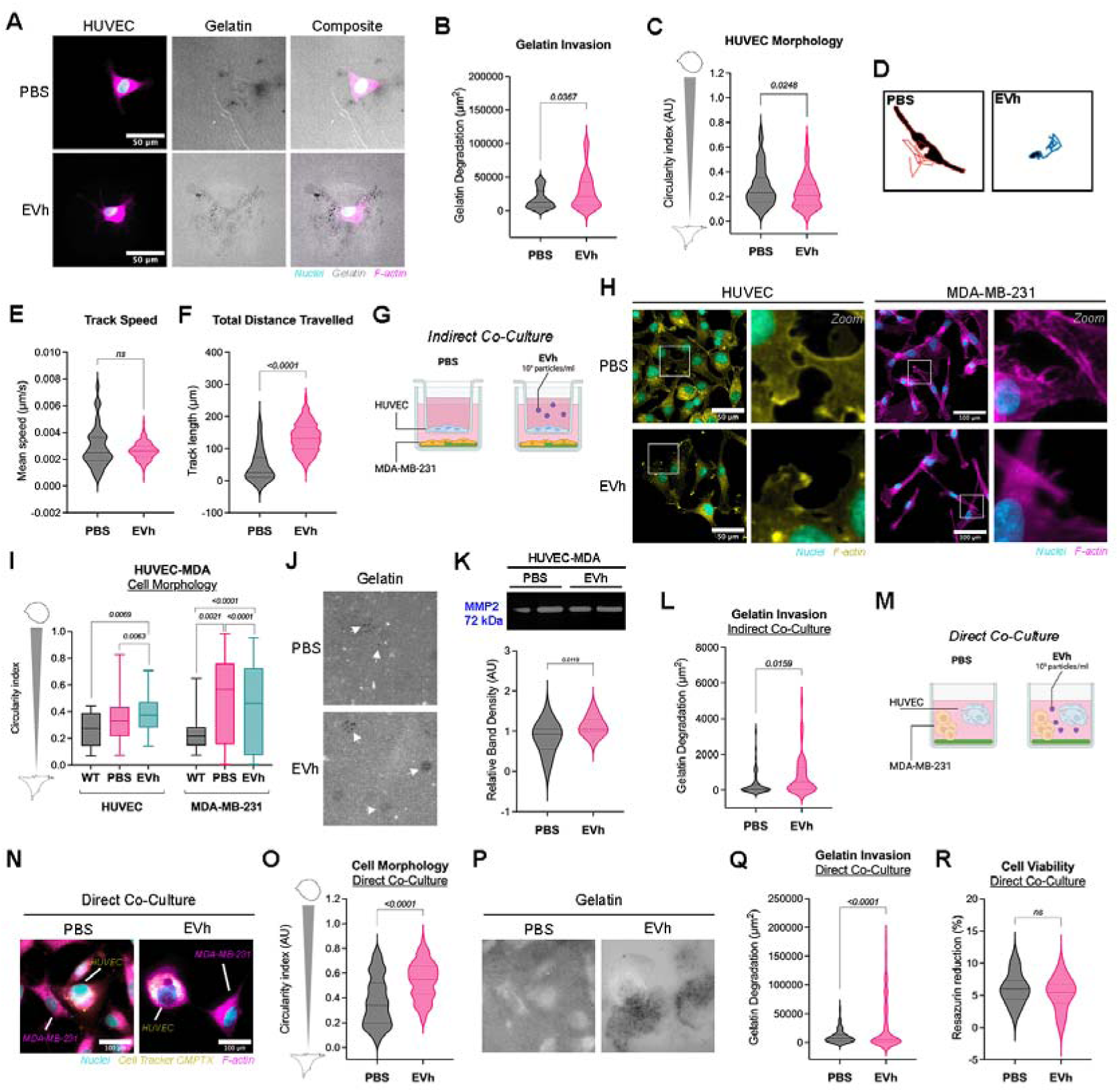
Endothelial cells respond to tumoral signaling. **A -** Representative images of HUVEC (nuclei in cyan, F-actin in magenta) treated with PBS or EVh on a gelatin coating (gray), with degradation spots in black (scale bar: 50 µm). **B –** Quantification of degraded gelatin area (µm^2^) in individual HUVEC cultures. **C –** HUVEC circularity index in individual culture. **D -** Tracking of HUVEC trajectory over 24h. **E –** Average track speed of cells in 24 h (µm/s). **F -** Total distance traveled by cells in 24 h (µm). **G –** Experimental design of the indirect co-culture assay. **H –** Representative images of HUVEC (nuclei in cyan, F-actin in yellow; scale bar: 50 µm) in indirect co-culture with MDA-MB-231 (nuclei in cyan, F-actin in magenta; scale bar: 100 µm). **I –** Cell circularity index for quantification of HUVEC and MDA-MB-231 morphology in direct co-culture. **J –** Representative images of gelatin matrix (gray) with degradation spots identified by white arrows. **K -** MMP-2 detection by gelatin zymography and densitometry analysis of indirect co-culture. **L –** Quantification of degraded gelatin area (µm^2^) in indirect co-culture. **M -** Experimental design of the direct co-culture assay. **N –** Representative images of HUVEC (cytoplasm in yellow) in direct co-culture with MDA-MB-231 (nuclei in cyan, F-actin in magenta) seeded atop a gelatin coating (gray) (scale bar: 100 µm). **O -** Cell circularity index for quantification of combined cell morphology in direct co-culture. **P -** Quantification of degraded gelatin area (µm^2^) in direct co-culture. **Q –** Cell viability of HUVEC and MDA-MB-231 direct co-culture. Assays were repeated on three different occasions with technical triplicates. *p* values indicated above comparative bars with statistical significance.

To explore a more complex level of tumoral communication, the pro-tumoral effect of EVh in endothelial cells was further investigated using indirect and direct co-culture assays with MDA-MB-231 cells. In indirect co-culture, where tumor cells were separated from endothelial cells by a porous membrane (Fig. 3G), HUVEC exhibited typical morphology and significant intercellular interaction, while the tumoral cells showed multiple cytoskeleton extensions, which gave them a downy appearance (Fig. 3H). In EVh-treated conditions, the cytoskeleton retracted as endothelial cells visibly lost intercellular interactions, and tumor cells showed a more elongated phenotype, without F-actin extensions (Fig. 3I). Tumoral gelatin invasion was detected in both co-culture groups, likely due to MMP-2 activation, which was detected only in its active form (Fig. 3J-K; Suppl. Fig. 2B). Both MMP-2 activity and gelatin degraded area were slightly higher in EVh-treated groups (Fig. 3L). Adding to previous results (7), we found EVh directs movement in MDA-MB-231, which facilitates tumoral invasion in gelatin matrices (Fig. Suppl. 2C-E).

In the direct co-culture approach, where both cell types were seeded in the same well simultaneously (Fig. 3M), close proximity triggered a stress response characterized by a star-shaped rearrangement of the endothelial cytoskeleton, while tumor cells appeared more cohesive (Fig. 3N). Upon treatment with EVh, cells regained their original morphological features (Fig. 3N; Suppl. Fig. 2F,G). These alterations were evident in the circularity index, highlighting the role of EVh in modulating intercellular interactions within the TME (Fig. 3O). Similar to the indirect co-culture model, EVh-treated cells also showed enhanced gelatin invasion (Fig. 3P–Q). Under these conditions, cell viability was unaffected (Fig. 3R).

### EVh modulates fibroblast behavior and suppresses tumor-associated invasion

Fibroblasts play a pivotal role in the tumor microenvironment by supporting tumor progression and remodeling the extracellular matrix. In this context, HDFa cells exhibited pronounced gelatinase activity, which was further enhanced following EVh treatment (Fig. 4A). This was accompanied by a migratory phenotype characterized by increased gelatin matrix degradation and a significant elevation in cell viability upon EVh administration (Fig. 4B–D). EVh also induced a polarized migration pattern in HDFa, reflected by decreased track speed and increased total distance traveled (Fig. 4E–G).

**Figure 4.**
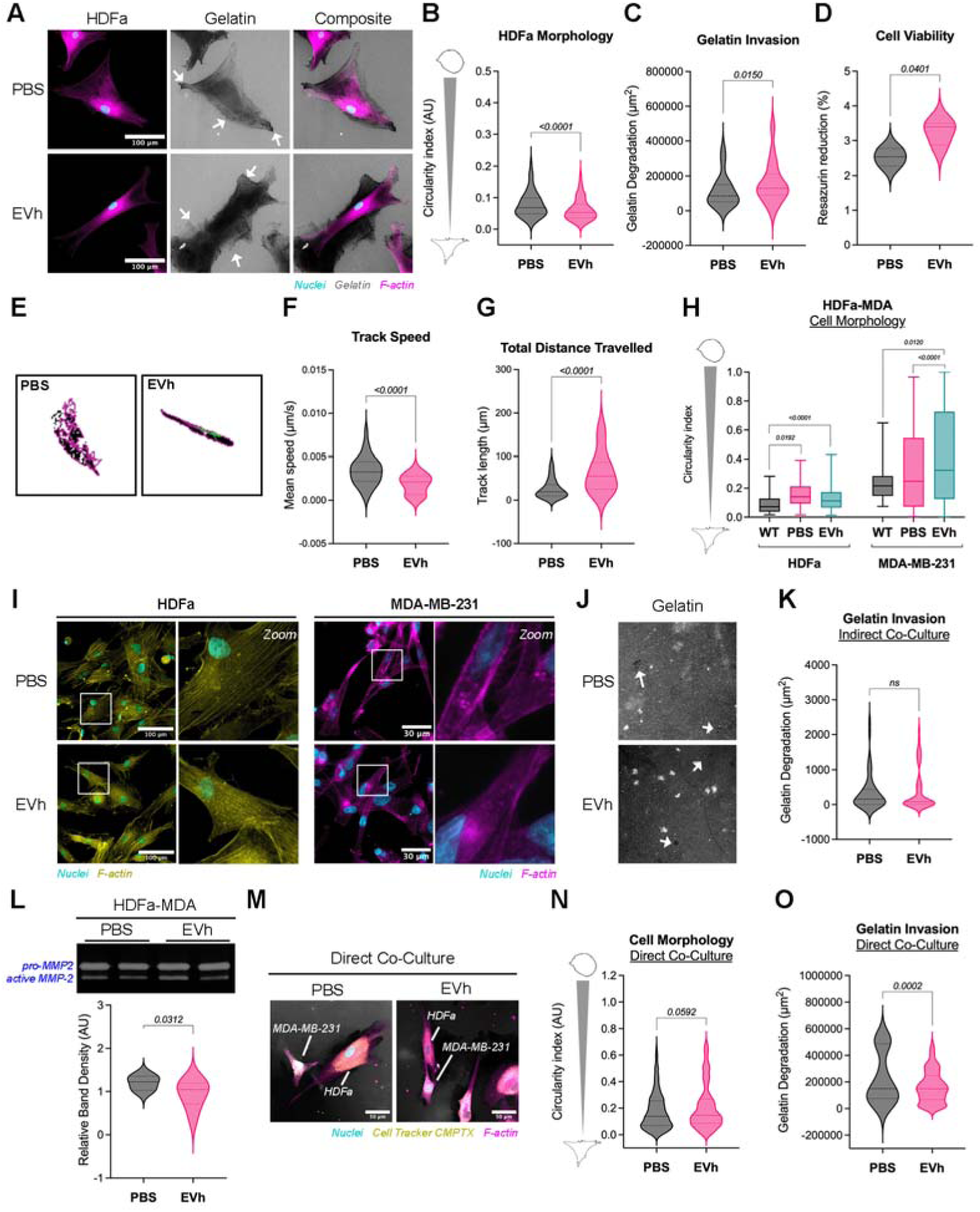
Fibroblast response to tumoral signaling. **A -** Representative images of HDFa (nuclei in cyan, F-actin in magenta) treated with PBS or EVh on a gelatin coating (gray), with degradation spots identified by white arrows (scale bar: 100 µm). **B –** Cell circularity index for quantification of HDFa morphology in individual culture. **C -** Quantification of degraded gelatin area (µm^2^) in individual HUVEC cultures. **D –** HDFa cell viability in individual culture. **E -** Tracking of HDFa trajectory over 24h. **F –** Average track speed of cells in 24 h (µm/s). **G -** Total distance traveled by cells in 24 h (µm). **H –** Representative images of HDFa (nuclei in cyan, F-actin in yellow; scale bar: 100 µm) in indirect co-culture with MDA-MB-231 (nuclei in cyan, F-actin in magenta; scale bar: 30 µm). **I –** Cell circularity index for quantification of HDFa and MDA-MB-231 morphology in indirect co-culture. **J -** Representative images of gelatin matrix (gray) with degradation spots identified by white arrows. **K -** Quantification of degraded gelatin area (µm^2^) in indirect co-culture. **L –** MMP-2 detected by gelatin zymography and densitometry analysis to identify gelatinase activity in indirect co-culture. **M –** Representative images of MDA-MB-231 (cytoplasm in yellow) in direct co-culture with HDFa (nuclei in cyan, F-actin in magenta) seeded atop a gelatin coating (gray) (scale bar: 100 µm). **N -** Cell circularity index for quantification of combined cell morphology in direct co-culture. **O –** Quantification of degraded gelatin area (µm^2^) in direct co-culture. Assays were repeated on three different occasions with technical triplicates. *p* values indicated above comparative bars with statistical significance.

To further assess fibroblast responses in distinct tumor-associated contexts, an indirect co-culture assay was established between HDFa and gelatin-precoated MDA-MB-231 cells. In this setup, fibroblasts displayed only modest alterations in response to tumor-derived cues, regardless of EVh treatment (Fig. 4H–I). A reduction in cell area was observed, leading to a broader morphology compared to monoculture controls, while gelatin degradation by tumor cells remained largely unaltered (Fig. 4J–K). These morphological changes were supported by a decreased circularity index. Zymographic analysis indicated that EVh treatment reduced MMP-2 activity in this context (Fig. 4L).

In a direct co-culture model, EVh appeared to suppress pro-invasive characteristics. Both fibroblasts and tumor cells displayed distinct morphological alterations (Fig. 4M, Suppl. Fig. 2H), with increased cellular circularity (Fig. 4N) and reduced gelatinase activity following EVh exposure (Fig. 4O). Notably, cell viability was unaffected under these conditions (Suppl. Fig. 2I).

### EVh attenuates phagoptosis and promotes pro-tumoral features in a multicellular circulating co-culture model

A multicellular circulating co-culture (MC-CC) was developed to recreate the cellular complexity of a TNBC tumor *in vitro* for evaluating EVh response in a TME-like setting (Fig. 5A). In the system, each TME cell type was seeded in a compartment for individual assessment of targets, while maintaining communication by a constant flow of 50 µl/s, set by previous optimization steps containing isolated cells (Fig. Suppl. 3A). A closed system comprised of MDA-MB-231, HUVEC, HDFa and THP-1 cells circulated for 24 hours showed high levels of phagoptosis (i.e., death by macrophage phagocytosis) (25), especially in the TNBC and endothelial cell chambers (Fig. 5B). In these chambers, integer nuclei and cytoplasm were seldomly found. EVh exposure led to more frequently sighted cells with integer nuclei and cytoplasm throughout chambers, suggesting phagoptosis inhibition. Cell morphological plasticity was confirmed by the circularity indexes of MDA-MB-231 and HUVEC (Fig. Suppl. 3B-C). The media flow reduced gelatin invasion in TNBC cells compared to individual static controls. EVh treatment partially recovered the cell invasive ability (Fig. 5C). The conditioned media was tested for gelatinases, and MMP-2 activity remained unchanged (Fig. 5D, Suppl. 3D). Cytokine IL-1β was present in similar levels in both groups, but IL-6 was more abundant in the EVh-treated system (Fig. 5E,F).

**Figure 5.**
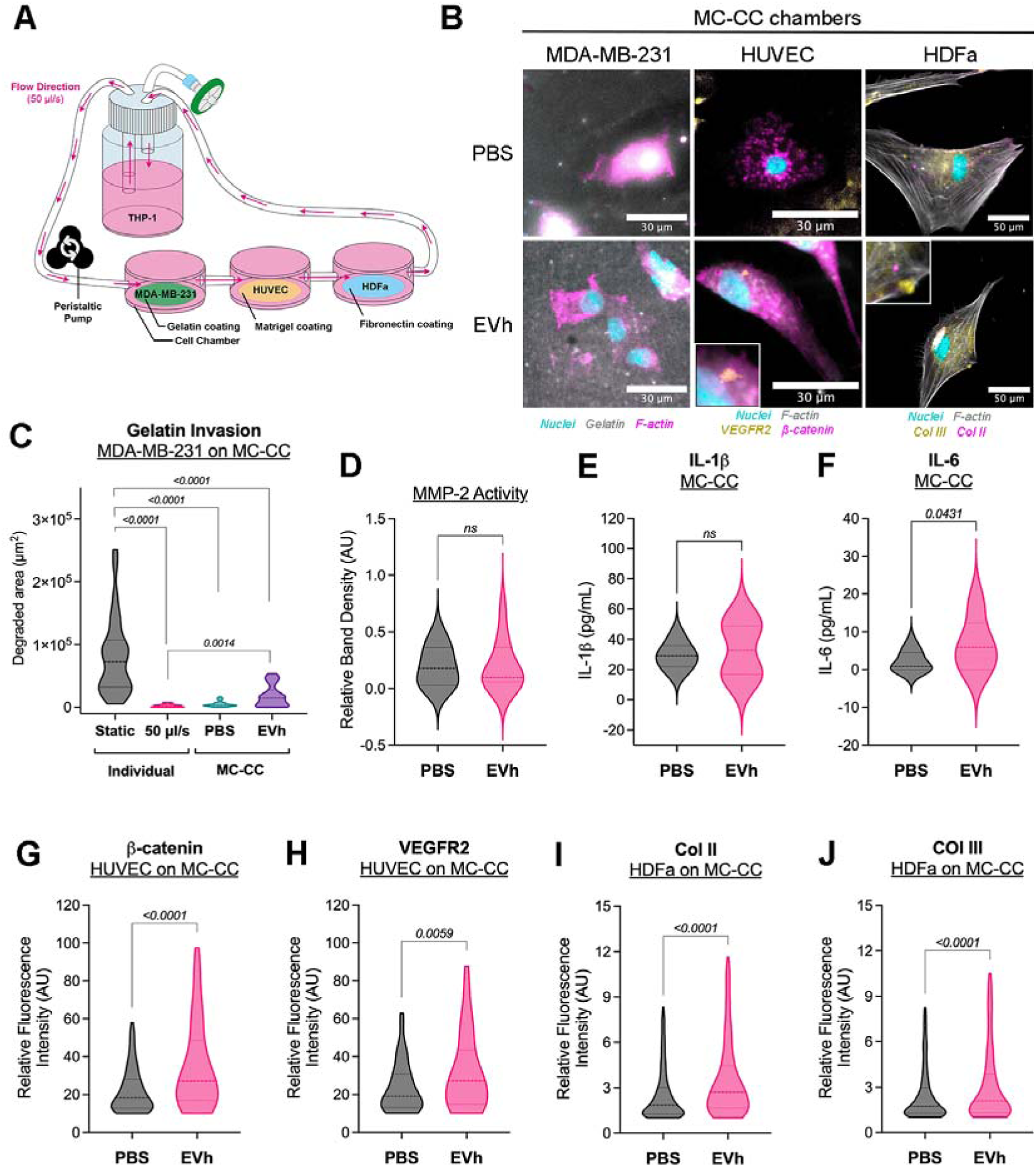
EVh inhibits phagoptosis in the TME. **A –** Experimental design of the multicellular circulating co-culture assay (MC-CC) using the QV500 system (Kirstall). **B -** Representative images of MC-CC chambers containing MDA-MB-231, HUVEC or HDFa treated with PBS or EVh. (scale bars: 30 µm and 50 µm). **C -** Quantification of degraded gelatin area (µm^2^) in the MC-CC MDA-MB-231 chamber. **D –** Band densitometry of MMP-2 detected by gelatin zymography from the MC-CC conditioned media. **E,F –** Cytokines IL-1β and IL-6 probed from the MC-CC conditioned media by ELISA. **G,H –** Relative fluorescence intensity of β-catenin and VEGFR2 in HUVEC, respectively. **I,J –** Relative fluorescence intensity of collagen II and III in HDFa, respectively. Assays were repeated on four different occasions with a single replicate. *p* values indicated above comparative bars with statistical significance.

Pro-tumoral features were assessed through quantitative immunofluorescence analysis. In endothelial cells, the expression of β-catenin and VEGFR2—key mediators of intercellular adhesion and the angiogenic cascade, respectively—was upregulated in the EVh-treated group (Fig. 5G,H). Among the cell types evaluated, fibroblasts exhibited the highest degree of structural preservation, with only minor morphological alterations detected (Fig. Suppl. 4E). Regarding extracellular matrix (ECM) remodeling, EVh exposure led to increased expression of collagen types II and III, predominantly localized around the pericellular region (Fig. 5I,J). Due to technical limitations, collagen type I expression was not analyzed.

Macrophage distribution within the MC-CC model revealed a predominant localization of THP-1 cells in the MDA-MB-231 chamber (Fig. 6A). Notably, the frequency of macrophage presence was reduced in EVh-treated conditions compared to controls, supporting the hypothesis that EVh mitigates phagoptosis within the tumor microenvironment. To further validate this observation, direct co-culture assays between THP-1 cells and each individual cell type were conducted. These experiments recapitulated the MC-CC findings, with elevated levels of phagoptosis observed in MDA-MB-231 and HUVEC cultures, and evidence of cellular debris present in fibroblast cultures (Fig. 6B). EVh treatment attenuated macrophage phagocytic activity and preserved the original morphology of all cell types. Interestingly, THP-1 cells formed a peripheral layer around fibroblasts, indicative of potential bidirectional communication. In this condition, fibroblasts acquired a more branched, polarized morphology, consistent with an activated state (Fig. 6C). Cell viability was unaffected by EVh across all co-culture settings (Fig. Suppl. 3F). However, EVh treatment enhanced gelatin invasion in tumor and endothelial cells following co-culture with THP-1, while fibroblast invasiveness was reduced (Fig. 6D).

**Figure 6.**
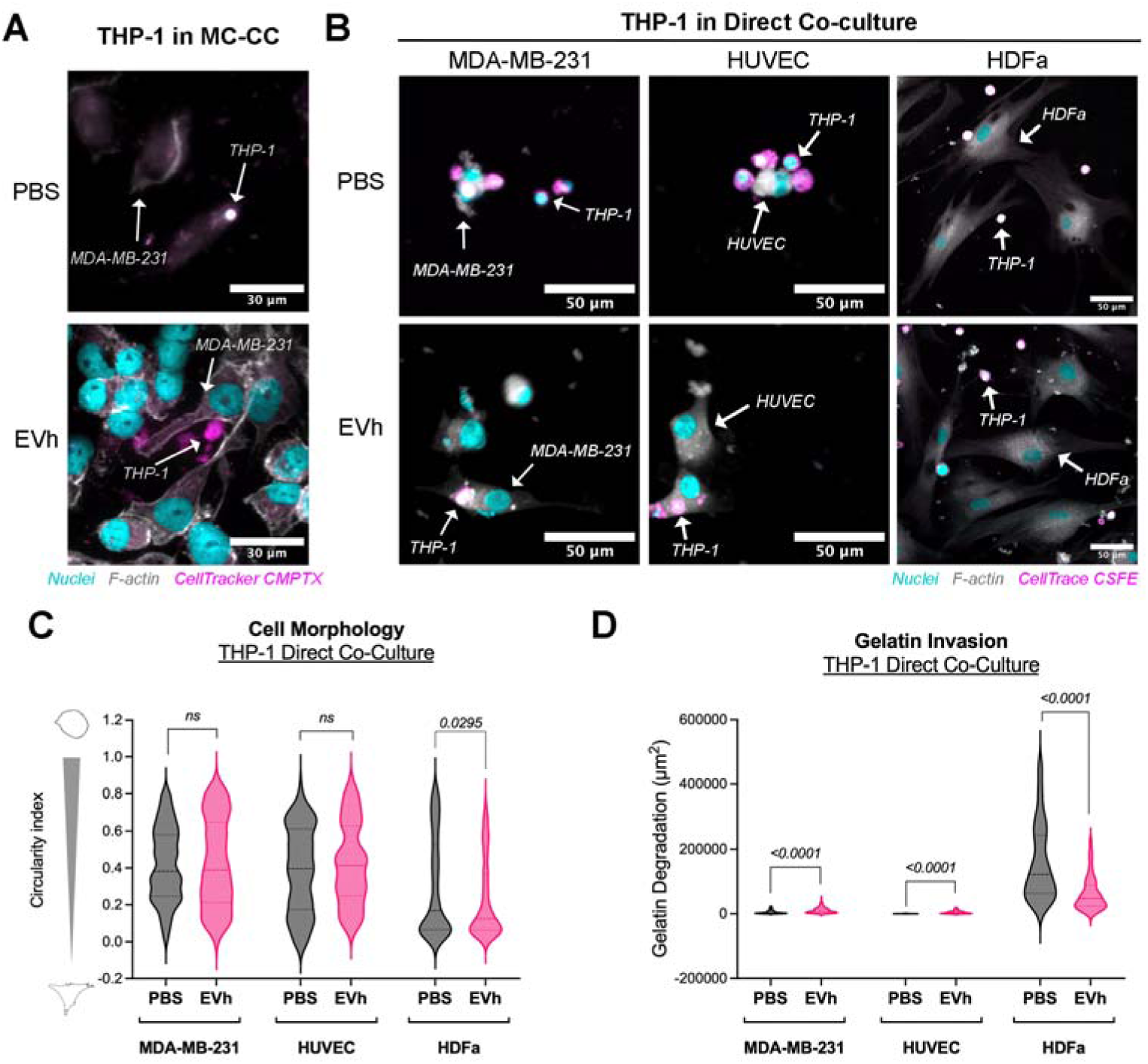
Macrophage remodeling by EVh in TME models *in vitro*. **A** – Representative images of THP-1 (magenta, stained with CellTracker CMPTX) attachment to MDA-MB-231 MC-CC chambers (nuclei in cyan, F-actin in gray; scale bar: 30 µm). **B -** Representative images of THP-1 cells (magenta, stained with CellTrace CSFE) in direct co-culture with MDA-MB-231, HUVEC or HDFa (nuclei in cyan, F-actin in gray; scale bars: 50 µm). **C -** Cell circularity index for quantification of combined cell morphology in THP-1 direct co-culture with MDA-MB-231, HUVEC and HDFa. **D –** Quantification of degraded gelatin area (µm^2^) in THP-1 direct co-culture with MDA-MB-231, HUVEC and HDFa. Assays were repeated on three different occasions with technical triplicates. *p* values indicated above comparative bars with statistical significance.

## Discussion

The role of EVs in tumor progression and development is well described, and as new facets of tumorigenesis are revealed, their importance to cancer continuity becomes apparent.

Tumor-associated EVs have been implicated in promoting angiogenesis (26), immune modulation (27), tumoral invasion (7,28), migration (29) and organotropic metastasis (30). The establishment of hypoxia in tumor progression promotes EV secretion in breast cancer, with distinct composition and characteristics (5–7). After describing the pro-invasive potential of hypoxic EV in comparison to normoxic EVs from TNBC cell line MDA-MB-231 (7), we show in this work that EVh differentiated THP-1 cells in M2-like macrophages, downregulating phagoptosis in tumor microenvironment models *in vitro*. Moreover, under the MC-CC model, EVh protected endothelial and tumoral cellular integrity, promoted pro- tumoral responses in endothelial cells and induced collagen synthesis in fibroblasts.

Monocyte recruitment and activation are crucial steps of the tumor-induced immune cascade, often triggered by the macrophage colony-stimulating factor (M-CSF) and other environmental cues (31). The differentiation of monocytes to macrophages *in vitro* relies on PMA to generate non-polarized cells, which may be induced to a pro-inflammatory M1 phenotype or an anti-inflammatory M2 profile, each with a distinct cytokine secretion pattern (17,32). We found that PMA-induced cells were enriched with pro-inflammatory components IL-1β, IL-6 and TNF-C, suggesting an M1-like profile, and the subtle reduction in IL-6 and sustained expression of anti-inflammatory IL-10 suggested that EVh promoted an M2-like phenotype (32–35).

The macrophage profiles were consistently maintained during co-culture with tumoral and endothelial cells, where phagoptosis was abundant in the control groups but avoided with EVh treatment. Cell death by phagocytosis results from the imbalance between the eat me and don’t eat me signals from the target cell and their recognition by receptors in the phagocytic cell (25). In our models, EVh likely modulated don’t eat me signals, such as CD47 and MHC-I, to improve cell survival, corroborating previous findings in which hypoxia inhibited macrophage phagocytosis in a HIF-1C-dependent manner (36,37). Further studies are required to confirm this hypothesis.

Studying the TME *in vitro* is challenging due to methodological, practical, and technological limitations. The recent development of techniques such as multicellular co-cultures, microfluidics and 3D culture has improved the complexity with which we can investigate basic mechanisms (38–40). A significant limitation of multicellular 3D co-cultures is the difficulty of evaluating responses in individual cells from the cultured mass, which often relies upon more refined and less accessible techniques (15). To circumvent this issue, we described a novel method to study the TME *in vitro* where cells are cultured *in tandem* in a commercially available system, henceforth named a multicellular circulating co-culture (MC-CC). Results obtained from the MC-CC model were complementary to other co-culture techniques, including direct and indirect co-culture, but we have not investigated whether it is a better alternative to 3D culture. This method was previously described for studying the blood-brain barrier but has several limitations, including the number of available co-culture chambers, flow shear force, and lack of direct contact between chambers which reduces our ability to evaluate certain cellular responses (41). We acknowledge that different experimental designs, including other cell types, order of chambers, incubation conditions, flow rate, and media composition, may interfere with the results and have not been investigated in the present study.

Our experimental design permitted the evaluation of cell morphology alongside cytokine and enzyme secreted in the circulating medium across all MC-CC chambers. EVh cause stromal and tumor cells to favor a more pro-tumoral phenotype, likely due to the increased levels of IL-6, a cytokine whose overexpression is closely related to the development of various tumors, regulation of the acute phase of inflammation, and modulation of T and B lymphocytes (42). Furthermore, the high levels of IL-6 and IL-1β favor the inflammatory phenotype of cancer-associated fibroblasts, which is consistent with our evaluation of fibroblast response (43). There was a non-statistical tendency for an increase in IL-1β levels in EVh-treated cells, suggesting fibroblast activation, but additional testing to detect differences in C-smooth muscle actin (C-SMA) is required to confirm this supposition.

EVh in endothelial cells under MC-CC increased the expression of markers for cell-cell adhesion and proangiogenicity, including VEGFR2 and β-catenin (44). This may facilitate tumor development and metastatic spread by increasing the connections between endothelial and tumoral cells and encouraging angiogenesis (44,45). Intercellular junctions are crucial for endothelial integrity and depend on the function of multiple proteins, especially VE-cadherin (46). With nearly exclusive expression in vascular endothelium, VE-cadherin is intracellularly linked to β-catenin, which directly connects to the cytoskeleton (47,48). In the indirect co-culture assay, we could not identify which adhesive molecules were influenced by EVh; however, multicellular co-culture under flow suggested an increase in β-catenin expression in HUVEC under EVh action. One possible interpretation for the increased β-catenin is that, since HUVECs are not fully confluent and VE-cadherin localization at the cell membrane is not evident, β-catenin may be predominantly cytoplasmic or even translocated to the nucleus, where it can promote transcriptional programs associated with cell migration rather than strengthening cell-cell junctions. This scenario aligns with the observed increase in VEGFR2 expression and suggests that β-catenin upregulation may contribute to enhanced endothelial migration and angiogenic activity rather than contradicting the pro-adhesive environment. These findings underscore the complexity of endothelial responses within the tumor microenvironment and warrant further investigation. Furthermore, we suggest that EVh disrupts endothelial cell adhesion to aid EVh-mediated tumor cell invasion.

Fibroblasts are structural support cells that can associate with tumors, transforming into cancer-associated fibroblasts (CAFs) through prolonged tumor-associated inflammation (TAI) signaling (43). TAI is sustained by continuous tumor mass growth and hypoxia, which have been described as determinants of the inflammatory phenotype of fibroblasts in pancreatic cancer (49). EVh increased fibroblast activity, modified their morphology and, in MC-CC, promoted the synthesis of type II and type III collagen, Both collagens are fibrillar and associated with increased ECM stiffness, a crucial factor for metastasis (50). Hypoxic extracellular vesicles also play a significant role in modulating fibroblast activity, leading to increased production of collagen types II and III. This process of extracellular matrix remodeling through MMP production and excessive collagen formation has been shown to promote tumor growth, progression, and resistance to treatments such as radiotherapy (44,45,51,52). In this setting, we show that hypoxic extracellular vesicles from triple-negative breast cancer impact fibroblast activity.

## Conclusion

In summary, we found that tumor-derived hypoxic EVs show a distinct immune-modulating function, allowing the differentiation of M2-like macrophages in a TNBC in the TME *in vitro* and inhibiting endothelial and tumoral phagoptosis. EVh administration protected the endothelial and tumoral cellular integrity from phagoptosis. It promoted pro-tumoral and pro-angiogenic responses in endothelial cells, inducing collagen synthesis in fibroblasts and their potential differentiation to CAFs. In addition, our work describes a novel method for studying the TME *in vitro* using a commercial system with great potential for investigating other tumor-related processes, including circulating tumor cells and metastasis.

## Author’s contribution

BCP: Conceptualization; Data curation; Formal Analysis; Funding acquisition; Investigation; Methodology; Project administration; Software; Writing – original draft. PHTB: Formal Analysis; Investigation; Visualization; AMM: Formal Analysis; Investigation; Writing – review & editing; CAC: Formal Analysis; Investigation; Methodology; Writing – review & editing; GG: Investigation; Visualization; LTG: Methodology; Visualization; MMG: Investigation; Writing – review & editing; ADZ: Investigation; Visualization; AMF: Conceptualization; Resources; Writing – review & editing; MRC: Funding acquisition; Resources; Software; Writing – review & editing; WFA: Conceptualization; Resources; Supervision; Writing – review & editing; HSSA: Conceptualization; Funding acquisition; Resources; Supervision; Writing – review & editing.

## Declarations

The authors (BCP, PHTB, AMM, CAC, GG, LTG, MMG, ADZ, AMF, MRC, WFA, HSSA) declare this study complies with the highest ethical guidelines and biosafety regulations. This study does not involve clinical or pre-clinical samples and did not require consent to participate. All authors declare no competing interests in this study and consent to publication. This study does not contain artificial intelligence-generated data.

## Supporting information

Suppl.Inf.

## Acknowledgments

The authors thank members from LMBBM and LABEN for critical advice. We thank the Laboratory of Structural Characterization (LCE/DEMa/UFSCar) for the electron microscopy facilities; Otavio Thiemann and Glaucius Oliva (IFSC/USP) for Beckman ultracentrifuge and rotors; Fausto Bruno dos Reis Almeida for the use of Nanosight NS300; Monica Rosa da Costa Iemma for the HDFa cell lineage; Gerson Jhonatan Rodrigues and Luis Henrique Oliveira de Moraes for the HUVEC cell lineage.

## Data availability

The data generated in this study are available within the article and its supplementary data files. Raw files are available upon reasonable request to the corresponding author. Detailed protocols have been submitted to the “protocols.io” platform (20,21,23,24). We have submitted all relevant data of our EV experiments to the EV-TRACK knowledgebase (EV-TRACK ID: EV240149) (53).

## Funding

This work was funded by the São Paulo Research Foundation through grants 2019/05149-9, 2021/01983-4, and 2022/04146-9 (to BCP); 2022/12307-2 (to PHTB); 2020/11328-0 (to LTG); 2017/01287-2 (to AMF); 2021/01863-9 and 2021/14673-3 (to MRC); 2019/11437-7 (to HSSA); Coordenação de Aperfeiçoamento de Pessoal de Nível Superior (CAPES), code 001; and Conselho Nacional de Desenvolvimento Científico e Tecnológico (CNPq) through grants 143120/2023-9 (ID 1618, to GG); 174872/2023-2 (to AMF). The funding agencies had no direct involvement in the study’s design, conduct, analysis, or reporting.

## Notes

### Competing Interest Statement

The authors have declared no competing interest.

### Summary of Updates

The manuscript was thoroughly corrected to include the reviewer's considerations.

