## Supplementary material for "Tumoral Hypoxic Extracellular Vesicles Foster a Protective Microenvironment in Triple-Negative Breast Cancer": Suppl.Inf.

### Table of contents

Supplementary Figure 1: Timelapse of macrophage differentiation p. 2
Supplementary Figure 2: Additional HUVEC and HDFa responses to tumoral signaling p. 3
Supplementary Figure 3: Multicellular circulating co-culture p. 4

**Supplementary Figure 1. Timelapse of macrophage differentiation.**

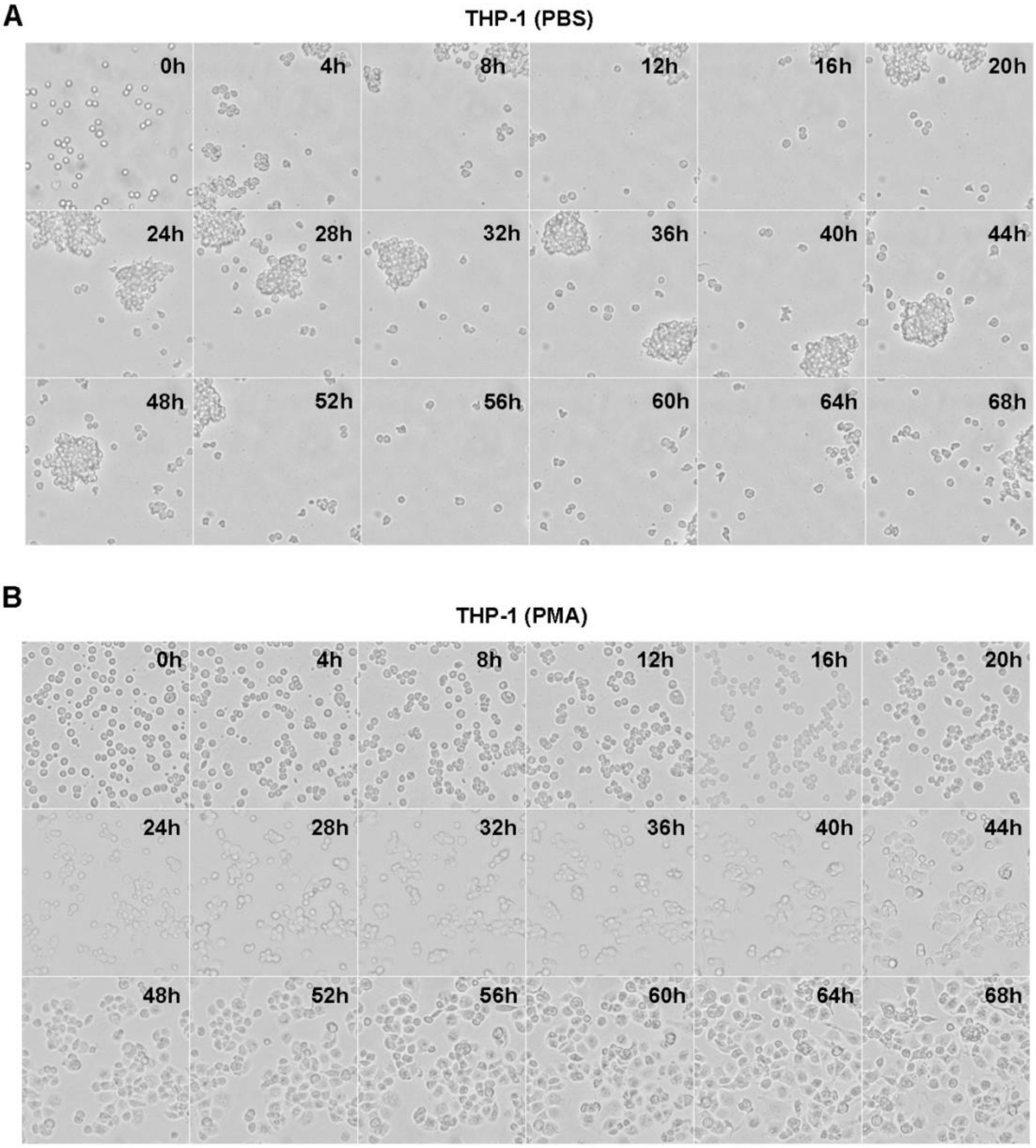


**A -** Tracking of untreated THP-1 cells (i.e., PBS) and **B –** THP-1 cells treated with PMA. Snaps from the same site were compiled by 4-hour increments over 3 days using the CytoSmart Lux 2 timelapse microscope (Lonza).

### Supplementary Figure 2: Additional HUVEC and HDFa responses to tumoral signaling

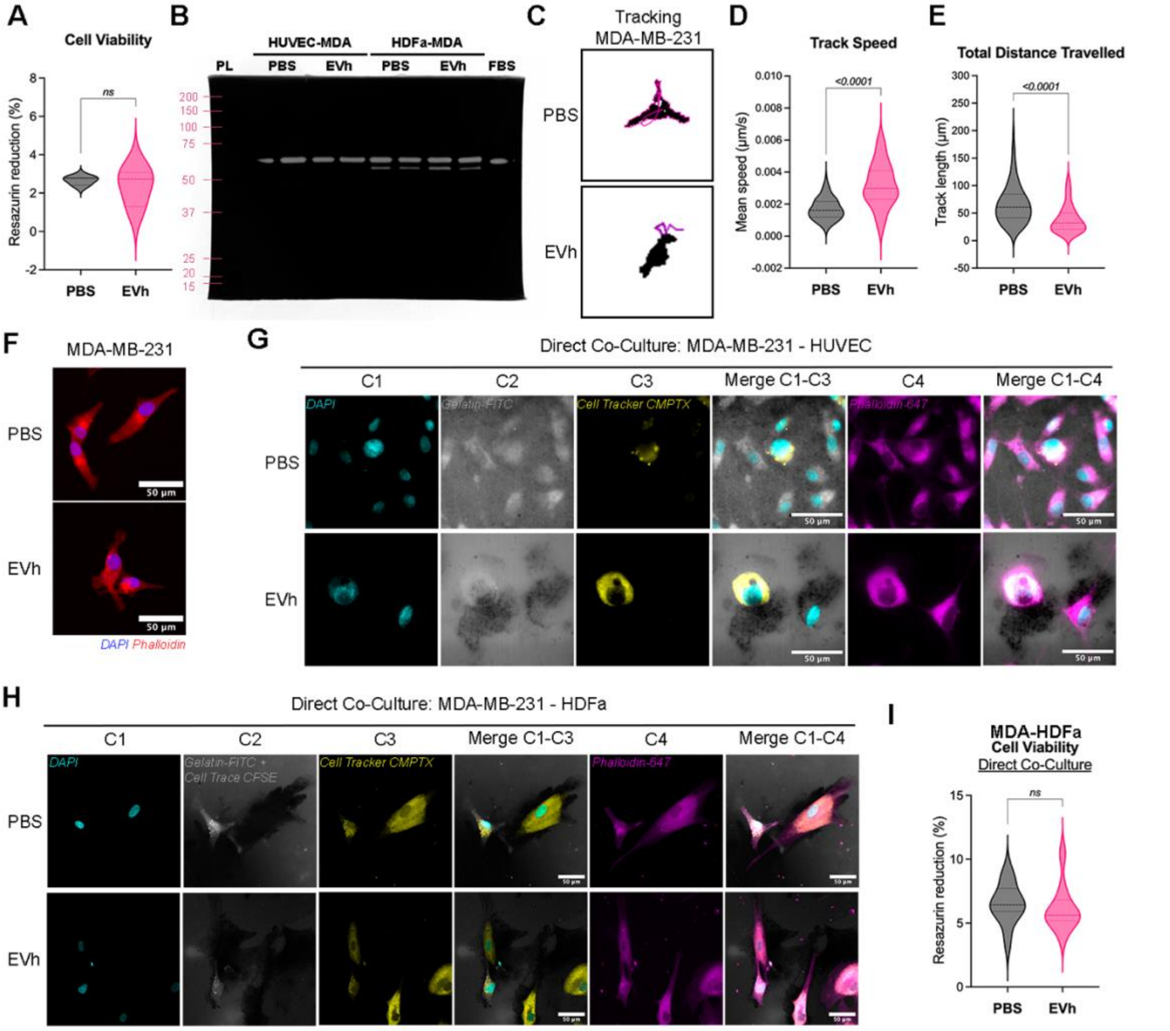


**A -** Cell viability of untreated (i.e., PBS) and EVh-treated individual HUVEC culture. **B –** Complete zymogram of indirect co-culture conditioned media. **C –** Tracking of MDA-MB-231 trajectory over 24h. **D –** Average track speed of cells in 24 h (μm/s). **E -** Total distance traveled by cells in 24 h (μm). Movement tracking of untreated (i.e., PBS) and EVh-treated MDA-MB-231 cells. **F –** Representative images of MDA-MB-231 cells in individual cultures, with F-actin in red and nuclei in blue. **G –** Segregated fluorescence channels of nuclei (C1, cyan), gelatin (C2, gray), Cell Tracker CMPTX (C3, yellow) and F-actin (C4, magenta) with composites from the direct co-culture of MDA-MB-231 and HUVEC. **H -** Segregated fluorescence channels of nuclei (C1, cyan), gelatin and Cell Trace CSFE (C2, gray), Cell Tracker CMPTX (C3, yellow) and F-actin (C4, magenta) with composites from the direct co-culture of MDA-MB-231 and HDFa. Scale bars: 50 μm. **I -** Cell viability of HDFa and MDA-MB-231 in direct co-culture.

### Supplementary Figure 3: Multicellular circulating co-culture

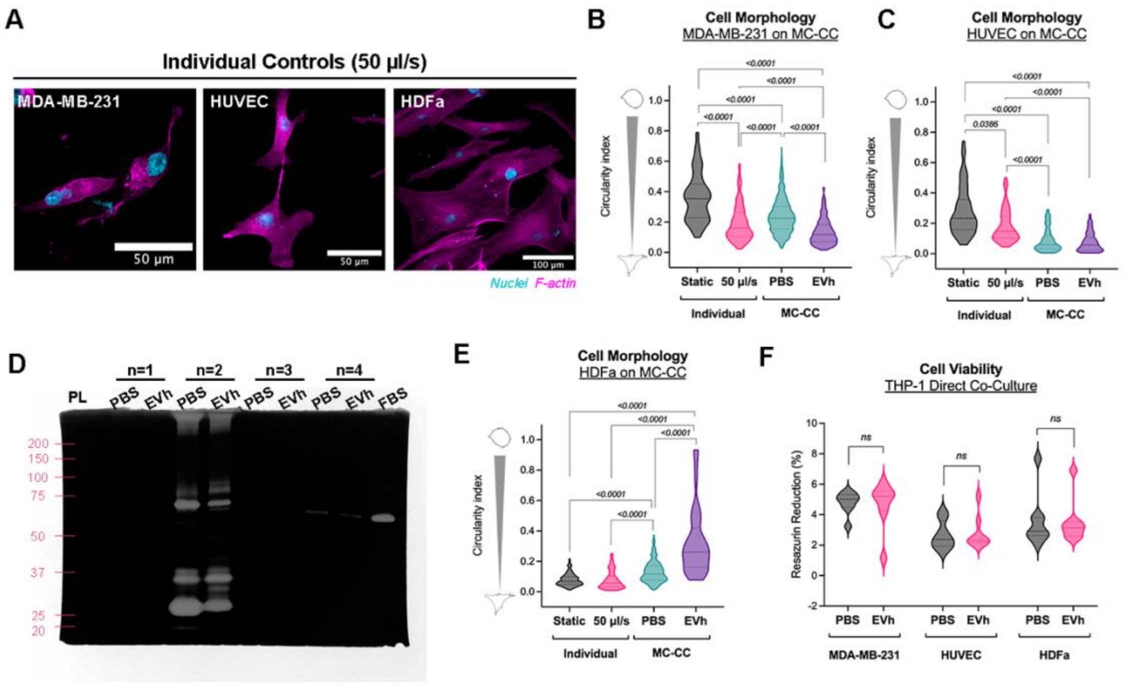


**A –** Individual circulating controls of MDA-MB-231, HUVEC and HDFa cells at 50 µl/s (nuclei in cyan, F-actin in magenta; scale bars: 50 µm and 100 µm). **B,C,E -** Cell circularity index for MDA-MB-231, HUVEC and HDFa in MC-CC, respectively, comparing untreated and EVh-treated systems with static and circulating individual controls. **D –** Complete zymogram of indirect co-culture. **F –** Cell viability of THP-1 direct co-culture with MDA-MB-231, HUVEC and HDFa. *p* values indicated above comparative bars with statistical significance.
